# The rise of the three-spined stickleback – eco-evolutionary consequences of a mesopredator release

**DOI:** 10.1101/2020.05.08.083873

**Authors:** Britas Klemens Eriksson, Casey Yanos, Sarah Bourlat, Serena Donadi, Michael C. Fontaine, Joakim P. Hansen, Eglė Jakubavičiūtė, Karine Kiragosyan, Martine E. Maan, Juha Merilä, Åsa N. Austin, Jens Olsson, Katrin Reiss, Göran Sundblad, Ulf Bergström, Johan S. Eklöf

**Author notes:** These authors jointly directed this work (Ulf Bergström, Johan S. Eklöf).

## Abstract

Declines of large predatory fish due to overexploitation are restructuring food webs across the globe. It is now becoming evident that restoring these altered food webs requires addressing not only ecological processes, but evolutionary ones as well, because human-induced rapid evolution may in turn affect ecological dynamics. In the central Baltic Sea, abundances of the mesopredatory fish, the three-spined stickleback (*Gasterosteus aculeatus*), have increased dramatically during the past decades. Time-series data covering 22 years show that this increase coincides with a decline in the number of juvenile perch (*Perca fluviatilis*), the most abundant predator of stickleback along the coast. We studied the interaction between evolutionary and ecological effects of this mesopredator take-over, by surveying the armour plate morphology of stickleback and the structure of the associated food web. First, we investigated the distribution of different stickleback phenotypes depending on predator abundances and benthic production; and described the stomach content of the stickleback phenotypes using metabarcoding. Second, we explored differences in the relation between different trophic levels and benthic production, between bays where the relative abundance of fish was dominated by stickleback or not; and compared this to previous cage-experiments to support causality of detected correlations. We found two distinct lateral armour plate phenotypes of stickleback, incompletely and completely plated. The proportion of incompletely plated individuals increased with increasing benthic production and decreasing abundances of adult perch. Stomach content analyses showed that the completely plated individuals had a stronger preference for invertebrate herbivores (amphipods) than the incompletely plated ones. In addition, predator dominance interacted with ecosystem production to determine food web structure and the propagation of a trophic cascade: with increasing production, biomass accumulated on the first (macroalgae) and third (stickleback) trophic levels in stickleback-dominated bays, but on the second trophic level (invertebrate herbivores) in perch-dominated bays. Since armour plates are defence structures favoured by natural selection in the presence of fish predators, the phenotype distribution suggest that a novel low-predation regime favours sticklebacks with less armour. Our results indicate that an interaction between evolutionary and ecological effects of the stickleback take-over has the potential to affect food web dynamics.

## Introduction

Human-induced biodiversity loss strikes particularly hard on higher trophic levels, and many populations of top predators have declined across the globe (Ripple et al. 2014). In marine systems, predator loss due to overfishing has altered trophic cascades, i.e. by inducing changes in predator-prey relations that propagate through food webs, affect biodiversity, and change ecosystem structure (Jackson et al. 2001, Daskalov 2002, Worm et al. 2006). In response to the severe economic and cultural losses resulting from these changes, various measures have been initiated to restore biodiversity at higher trophic levels (e.g. marine protected areas; Gell and Roberts 2003, Lester et al. 2009). However, these measures may be ineffective because of pervasive but poorly understood feedback mechanisms within the ecosystem (Nyström et al. 2012), as well as interactions between multiple human impacts that delay or prevent recovery (Frank et al. 2005, Eriksson et al. 2010, Frank et al. 2011).

Currently, we have an inadequate understanding of the drivers that structure biodiversity and food webs that are, themselves, changing. One such poorly understood driver is rapid evolution. Evolution was traditionally assumed to be slow and unimportant on ecological timescales, and has therefore rarely been included in ecological studies or ecosystem management (Fussman et al. 2007, De Meester et al. 2019, Hendry 2019). Today, however, we know that selection can change population dynamics on short timescales, and that overfishing in particular has the potential to cause exceptionally rapid evolution of target species (Conover and Munch 2002, Kuparinen and Merilä 2007, Hutchings and Fraser 2008, Audzijonyte et al. 2013). Over the past decade, we have also become increasingly aware that rapid evolution can feed back on to ecosystem properties, so called eco-evolutionary feedback (Post and Palkovacs 2009, Schoener 2011, Turcotte et al. 2011, Hendry 2017, Govaert et al. 2019). Still, evolutionary consequences of large-scale predator loss for lower trophic levels are poorly understood.

Declines of large predatory fish are typically followed by dramatic increases in the abundance of their main prey, often smaller fish species; a process known as “mesopredator release” (Prugh et al. 2009). For the prey fish, the strong increase in density intensifies intraspecific competition, and should in theory shift the selection regime from predation defence towards increased individual-level resource specialization and population-level resource use diversity (Svanbäck and Persson 2004, Svanbäck and Bolnick 2005, Svanbäck and Bolnick 2007). Accordingly, there are clear examples of how species populations have increased their competitive ability and diversified after escaping control from their native predators (Palkovacs et al. 2011, Mlynarek et al. 2017). Thus, mesopredator release caused by loss of large predators may lead to a strong selection pressure for different traits among prey fish. This may, in turn, impact other trophic levels by introducing new predator-prey interactions, ultimately impacting the nature and strength of trophic cascades.

Another human impact that interacts strongly with trophic cascades is increasing productivity e.g. trough nutrient addition to aquatic environments (e.g. Eriksson et al. 2009, Sieben et al. 2011b, Eriksson et al. 2012, Östman et al. 2016). Models of trophic dynamics suggest that predator effects on lower trophic levels interact with resource availability, such that the abundance of every second trophic level increases with primary production (Fretwell 1977, Oksanen et al. 1981, Nisbet et al. 1997, Oksanen and Oksanen 2000, Eriksson et al. 2012). In a system with four trophic levels the predator community accumulates biomass. Thus, for mesopredators (third trophic level), there should be an increase in selection for antipredator traits, but no increase in density, with increasing productivity (Oksanen and Oksanen 2000, Eriksson et al. 2012)(Fig. 1a). However, in a system with three trophic levels, energy (biomass) accumulates at the mesopredator level, while at the same time the absence of top predators result in a relaxed selection for antipredator traits. Here, increasing productivity would lead to an increase in mesopredator density, and consequently, density-dependent competition (Oksanen and Oksanen 2000, Svanbäck and Bolnick 2007, Eriksson et al. 2012). Thus, the number of trophic levels and the productivity of the ecosystem should together determine mesopredator abundance as well as trait distribution (Fig. 1a), which may in turn influence trophic dynamics through changes in, for example, diet composition, behaviour, and/or elemental stoichiometry; and thereby generate an eco-evolutionary feedback on food web structure (broad-sense eco-evolutionary feedback; Hendry 2017, de Meester et al. 2019)(Fig. 1b).

**Figure 1:**
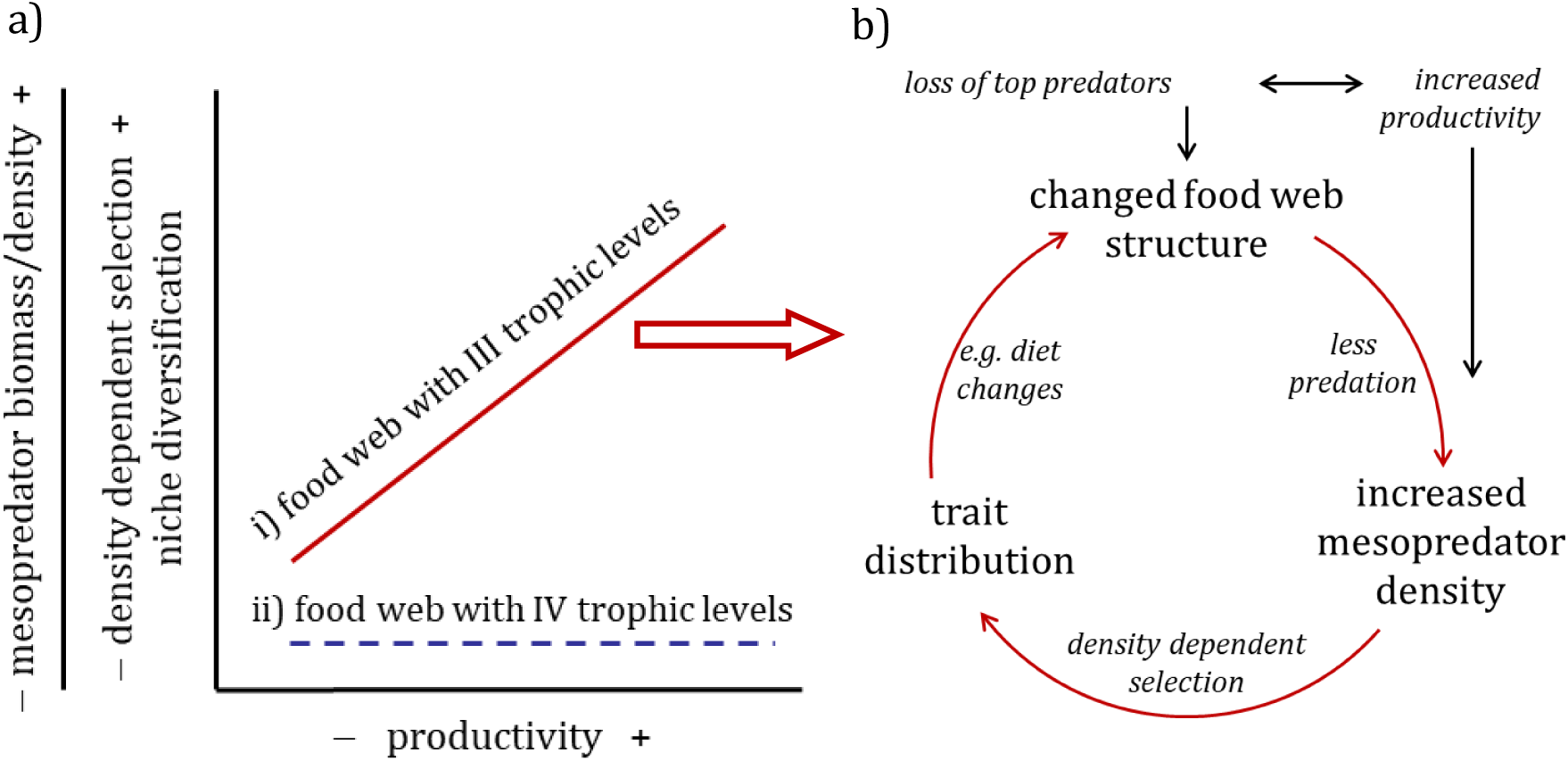
Conceptual model illustrating how the density development and trait distribution of a mesopredator population depend on the number of trophic levels and productivity of the system (according to the exploitation ecosystems hypothesis by Oksanen et al. 2000). With top predators, the system has four trophic levels (dotted blue line). Without top predators, the system has three effective trophic levels and mesopredators dominate the predator community (solid red line). a) In a system with three trophic levels, the mesopredator community accumulates biomass. Ecological effects: Increase in mesopredator density with increasing productivity; changed food web relations lead to a trophic cascade (Oksanen and Oksanen 2000, Eriksson et al. 2012). Evolutionary effects: the absence of top-predators results in a relaxed selection for antipredator traits; and increased selection for traits favourable under density-dependent competition for food (Svanbäck and Bolnick 2007). b) Conceptual model illustrating how increased mesopredator density and resulting changes in trait distribution (caused by the combination of predator loss and increased productivity) may influence trophic dynamics by changing in energy flow (e.g. changes in diet, elemental stoichiometry, behaviour), thereby leading to a broad-sense eco-evolutionary feedback on food web structure (Hendry 2017, de Meester et al. 2019).

In the western parts of the central Baltic Sea (NW Europe), densities of a small mesopredatory fish – the three-spined stickleback (*Gasterosteus aculeatus* L., hereafter stickleback) – have increased up to 45 times during the last decades, following regional and/or local declines of their natural predators; the large-bodied commercially and recreationally harvested fish cod (*Gadus morhua*), pike (*Esox lucius*), and perch (*Perca fluviatilis*; Ljunggren et al. 2010, Eriksson et al. 2011, Bergström et al. 2015, Olsson 2019, Olsson et al. 2019). Suggested mechanisms behind the stickleback population explosion include a combination of human impacts that link drivers of change across large spatial scales. Juvenile stickleback migrate offshore during autumn and, when mature, return to the coast for spawning in spring (Bergström et al. 2015). Offshore, overfishing of cod, increased water temperatures, and increased food availability due to eutrophication may have contributed to their increased survival (Eriksson et al. 2011, Lefebure et al. 2014). Inshore, local decreases in perch and pike populations may have increased recruitment and juvenile survival of stickleback in some areas (Östman et al. 2014, Bergström et al. 2015, Hansson et al. 2017). The local decreases in pike and perch are caused by loss or degradation of reproduction habitats due to wetland drainage, coastal constructions, and boating (Nilsson et al. 2014, Sundblad and Bergstrom 2014, Hansen et al. 2019) and, potentially, increased predation from seals and cormorants (Östman et al. 2014, Hansson et al. 2017). Today, there is a strong inverse relationship between local abundances of stickleback on the one hand and perch and pike on the other (Bergström et al. 2015). The dramatic increase in stickleback abundance, in combination with elevated resource availability from large-scale eutrophication, has caused a trophic cascade that has restructured near-shore food webs, increased filamentous algal blooms, threatening coastal biodiversity (Eriksson et al. 2009, Sieben et al. 2011a, Byström et al. 2015, Donadi et al. 2017).

Globally, the marine stickleback is known to have rapidly and repeatedly adapted to freshwater habitats with low predation pressure by evolving reduced armour phenotypes (i.e. lower number of bony lateral plates and loss/reduction of pelvic girdle), apparently in response to relaxed selection on these anti-predator traits (Bell 2001, Huntingford and Coyle 2006). A large body of research supports the hypothesis that fully plated marine individuals are better protected against predation by fish than individuals with reduced plating (e.g. Reimchen 1992, 2000). On the other hand, reduced lateral plate numbers can increase fast-start performance and manoeuvrability, and thereby provide means to avoid predation in habitats where refuge from predators (e.g. vegetation) is present (Bergström 2002, Leinonen et al. 2011b). In addition, adaptation to predation can strongly influence the feeding ecology of stickleback (reviewed in Bell and Foster 1994, Harmon et al. 2009).

The stickleback has three lateral plate phenotypes: fully plated individuals, which have plates along the sides of the body that run uninterrupted from the head to the tail fin (caudal end); partially plated individuals, which lack plates on the mid-section of the body; and low plated individuals, which only have plates on the head (pectoral) region of the body (Münzing 1963, Reimchen 2000). The plate phenotype is highly heritable and largely controlled by one major gene (Ectodysplasin; Colosimo et al. 2004), and hence, indicative of the underlying genotype (Loehr et al. 2012, DeFaveri et al. 2013). In the Baltic Sea, all three plate phenotypes of the three-spined sticklebacks have co-occurred for at least a century (Aneer 1974, Jakubavičiūtė et al. 2018), and there is evidence for adaptive phenotypic differentiation in number of lateral plates across the Baltic Sea (DeFaveri and Merilä 2013).

The aim of this study was to evaluate the possibility that the ecological and evolutionary effects of a mesopredator release have induced an eco-evolutionary feedback along the coast in the central Baltic Sea (as outlined in Fig. 1b). We first assessed temporal trends in the abundance of the three-spined stickleback and perch, the most common predator in the area, using time series data covering 22 years (1996-2017) to establish that there has been a shift in trophic structure and predation regime. Second, we assessed the consequences of the stickleback increase for ecological dynamics, using a field survey along a 360 km stretch of coast. We did so by investigating the relationship between potential production of the community and the accumulation of biomass on different trophic levels, comparing areas with different abundances of stickleback. To get an indication of the strength and causality of documented patterns, we compared the field data with results from a previous experiment in which the abundance of large predatory fish was manipulated (Sieben et al. 2011b). Third, we evaluated the possibility of eco-evolutionary feedbacks by relating differences in the spatial distribution of plate phenotypes to the abundance of predatory fish and potential production of the community. Finally, we documented differences in the diets of the different plate phenotypes, to evaluate the potential for feedbacks on food web structure caused by changes in plate phenotypes distribution.

We hypothesised (Fig. 1): 1) that in areas experiencing relatively high abundances of larger predatory fish, the local food web dynamics is characterized by four trophic levels; where the abundance of herbivores and potentially large predatory fish increases with production. 2) that in areas dominated by mesopredators, experiencing relatively high abundances of sticklebacks, the local food web dynamics is characterized by three trophic levels; where the abundance of sticklebacks and primary producers increases with production. 3) that in areas dominated by mesopredators, and thus with relatively low abundances of predatory fish, reduced selection for anti-predator traits results in an increased proportion of incompletely (low or partially) plated stickleback individuals at higher production (see Fig. 1a). 4) that the different stickleback plate phenotypes have different feeding ecology, as reflected by different prey composition in the stomach content.

## Material and Methods

### Trends in three-spined stickleback and juvenile perch abundances over time

We first analysed temporal trends in the abundance of juvenile three-spined stickleback and perch (the dominant coastal predator on stickleback). Data on temporal trends of juvenile three-spined stickleback and perch abundances from 1996 to 2017 was extracted from the Swedish national database for coastal fish (http://www.slu.se/kul). The data combined samplings of juvenile fish from various monitoring programs and research projects that quantified fish recruitment along the Swedish Baltic Sea coast. The available data on juvenile fish are spatially far more comprehensive than those for adults. Therefore, use of juvenile data enabled us to extract a time series that covered the full outer to inner archipelago gradient, which better represents the full range of habitats occupied by different species of fish than the regular coastal fish monitoring programs (see e.g. HELCOM 2018). The database on juvenile fish also includes small species, such as stickleback, which are poorly represented in the regular monitoring programs (as the multimesh gillnets used for monitoring lack mesh sizes small enough to capture stickleback in a representative way; HELCOM 2018).

We extracted abundances of juvenile perch and three-spined stickleback from an area covering a 360 km stretch of the central Baltic Sea coastline (Fig. S1.1). Fish were sampled in shallow bays using small underwater detonations; a standard method that catches fish with a swimbladder of lengths up to 20 cm within a blast radius of 2.5-5 m depending on detonation strength (1-10 g; size range of fish 2-10 cm; Snickars et al. 2007). Since fish are heterogeneously distributed at small spatial scales and the detonation method takes a snapshot of a defined area (ca. 20-80 m^2^ depending on detonation strength), the abundance data is highly variable. In addition, bays have been sampled at various intensities (from 5 to 90 detonations per bay, mean ± SD = 16 ± 11). We therefore square-root transformed abundance data from each detonation and then used averages for each bay for the statistical analyses. Moreover, we only included the years 1996 to 2017 in the analysis because before 1996 most years were only represented by one single bay; leaving 7553 detonations from 480 bay-year combinations in the dataset. The mean depth of all sampling was 1.7±0.59m (±SD) and nearly all sampling (94%) was conducted during late July to early September. All sampling was performed by certified personnel and with required ethical permits and exemptions from national fishing regulations.

We analysed trends in stickleback and perch abundances by fitting linear models with year as an explanatory variable. We did not perform a formal time series analysis accounting for autocorrelation of sampling stations over time, because most of the bays were sampled only once (184 out of 256 unique bays) and less than 10% were sampled more than four times. In addition, the replication of bays for each year was highly skewed: the first five years had an average replication of ca. 10 bays per year, compared to ca. 30 for the following decades. To balance the influence of the different time-periods on the model fit, we therefore used the averages for each year as response variable and treated each year average as a true replicate. However, this reduces the variability of the data, meaning that the R^2^ of the model fit only is related to the variation in yearly means; not to the variation between bays within or across years. Since the sub-set of bays were different each year, we included a spatial structure in the model to make sure trends were not caused by differences in sampled regions or characteristics of bays. The spatial structure consisted of adding average sampling depth, latitude, and longitude as explanatory factors to the model. We selected the best model for perch and stickleback abundances based on AICc (AIC for small samples); using the MuMIn package in R 3.5.2 (version 1.43.15; R Core Team, 2018.

Two years (1996 and 2000) were excluded from the model selection because they caused strong collinearity between latitude and longitude; sampling predominantly occurred in the southern and innermost part of archipelago. After the selection procedure, the omitted years were reintroduced into the final model. In the reduced dataset there was no correlation between the explanatory variables and we did not detect any problems with collinearity in the candidate models (the variance inflation factor (VIF): year=1.10, latitude=1.19, longitude=1.22, depth=1.06), or the final models (VIF: year =1.00, depth=1.00). If necessary, variables were normalized using square root transformations.

### Characterising the coastal food web and stickleback phenotype composition

Since the time series data described above mainly contain juvenile fish abundances, we characterized the benthic food web by sampling across trophic levels in 32 shallow coastal bays in spring 2014 (max sampling depth between 3 and 6 m). The area covered was the same as where the time-series data was extracted from (Fig. S1.1). The bays represented a gradient from inner to outer archipelago, characterized by gradients in wave exposure and bay morphology; factors that regulate the composition of the biological communities in the Baltic Sea archipelago (see Supplement 1 for detailed explanation of selection criteria and calculations). We characterized the physical environment of each bay by estimating wave exposure, distance to the open sea, and topographic openness (Ea). Wave exposure was estimated for each bay using a simplified wave model based on fetch, topography, and wind conditions using Geographical Information System (GIS) methods (Sundblad et al. 2014). Distance from the open sea was calculated by estimating the shortest water distance from the opening of each bay to the baseline (the starting point for defining territorial sea). Ea is defined as 100 × At/*a*: where At is the cross sectional area of the smallest section of the opening of the bay towards the sea and *a* is the bay surface area. Topographic openness is one of the most important factors structuring the coastal communities of Baltic Sea bays (Hansen et al. 2012, Hansen 2013). With increasing openness total-nitrogen decreases (since more isolated bays accumulate more nutrient-rich sediments) and spring-summer water temperature decreases (because shallow waters and smaller volumes warm more rapidly in isolated bays and high-water exchange of more open bays means larger influence of cold seawater) (e.g. Hansen 2013). This is reflected in higher phytoplankton and zooplankton biomass (Scheinin and Mattila 2010); and freshwater fish production in isolated bays (Snickars et al. 2009). However, with increasing openness the influence of seawater and the influx of nutrients from surrounding water bodies increases. This promotes the benthic community (Donadi et al. 2017), which is reflected in higher biomass of marine and brackish water algae and crustacean macro-herbivores (mainly benthic amphipods) in more open bays (Hansen et al. 2012, Hansen 2013). Thus, topographic openness is a good proxy for potential marine production in shallow Baltic Sea bays.

The fish community was sampled in spring (from May to early June) using three to six Nordic survey gillnets set out overnight in each bay (mesh size 5–55 mm, details in Supplementary Information, Supplement 1). All fish caught were identified to species, measured for total length, and counted. The number of individuals of each fish species caught were pooled for each bay and catch per unit effort (CPUE) was expressed as number of individuals per net and night of each species. In each bay, we estimated vegetation and filamentous algae cover, collected zooplankton, and measured temperature, turbidity, and fluorescence, at three to six randomly distributed stations (Supplement 1, Fig. S1.2a). Mesograzer (benthic macro-herbivores) biomass was quantified at each station by sampling epifauna in a randomly placed 0.2 × 0.2 m frame connected to a 1 mm-mesh bag (Supplement 1, Fig. S1.2b,c). After identification of mesograzers in the laboratory (i.e. shredders and gatherers, whose diet is dominated by macrophytes: see Donadi et al. 2017), the biomass of mesograzers was estimated as gram ash-free dry-mass (AFDM) using taxon-specific correlations between length and weight (Eklöf et al. 2017). The number of both stations and nets depended on the size of the bay. They were randomized within depth strata from a topological map to be representative of the depth profile in each bay, but restricted to a depth between 0.5 and 3 m (Supplement 1; also see Donadi et al. 2017 for a detailed description of sampling procedures).

In each bay, we sub-sampled 30 individuals of the sticklebacks caught in the nets for morphological and diet analyses; if ≤30 were caught, all individuals were included. To analyse body shape variation, eleven length parameters were documented from all individuals (following Jones et al. 2012, Supplement 2). The number of lateral plates were counted on both the left and right sides of the body and the average number of lateral plates for each individual was calculated. Plates were counted starting immediately after the operculum to the end of the caudal peduncle, following Aneer (1974). Plate phenotypes were identified by looking at gaps in the plate armour: low plated fish lacked caudal plates; partially plated fish had a clear gap between the main body plates and the caudal plates (the width of at least 2 body plates); fully plated fish had no clear gap in the plating. However, there were a number of phenotypes in the transition phase between low and partially plated individuals; for example, individuals with only a few clear body plates followed by a clear gap and then a keeled tail consisting of reduced plates. For the analyses, we therefore divided the phenotypes into completely (fully) or incompletely (low and partially) plated sticklebacks. In total, we counted plates of 560 individuals.

Algal production was estimated for each bay as percent cover of filamentous algae on settling plates following three months in the water (May to August). Each plate consisted of four 5 × 5 cm unglazed ceramic tiles glued in pairs to two bricks, which were placed flat on the bottom at ca 1.5 m depth (Fig. S1.2d). Algal cover on settling plates is a good measure of relative net primary production in the Baltic Sea (e.g. Eriksson et al. 2006, Eriksson et al. 2009, Eriksson et al. 2012), as filamentous algae respond to nutrient enrichment by fast accumulation of biomass, but are simultaneously grazed by invertebrate mesograzers (Worm et al. 2002, Råberg and Kautsky 2007). Mesograzers are in turn eaten by fish, and thereby play a key role in trophic cascades (Eriksson et al. 2009, Donadi et al. 2017). Consequently, the cover of filamentous algae is a good indicator of top-down (grazing) and bottom-up (nutrients) resource control of primary production. In eight of the 32 bays, individual tiles were either lost or completely/partially covered with sediment. These bays were therefore not included in the analysis of algae cover.

### Analysing food web structure

We tested the general hypothesis that three-spined stickleback abundances correlate with potential production and the abundance of predators along the coast. The CPUE of the three-spined stickleback in the 32 bays was analyzed by a General Linear Model that included the explanatory variables topographic openness (Ea) and CPUE of large perch, (number of individuals larger than 25 cm caught per net and night) using the lm function in R 3.5.2 (R Core Team, 2018). We used Ea as a proxy for net production of the marine benthic community (see above under “*Characterising the coastal food web*”; Donadi et al. 2017). Accordingly, a preliminary analysis showed that the vegetation cover in the bays correlated positively with Ea (r=0.51, n=32, p=0.003; Fig. S1.3). We used perch abundance as an indicator of predation pressure; perch is by far the most common stickleback predator in the coastal fish community. We only included perch >25 cm, as stickleback then constitute their most common prey item (>40 % of the diet in spring; Jacobson et al. 2019). The other common coastal fish predator, pike, is poorly represented by the gill-net catches because of their sedentary life-style, and was therefore not included in the analyses. CPUE data was normalized by square root transformations. There was no correlation between the explanatory variables topographic openness (Ea) and CPUE of large perch (Pearson’s product moment correlation: r=−0.19, t = −1.03, df = 30, p=0.309), and we did not detect any problems with collinearity in the model (VIF=1.04).

To get an indication of the number of effective trophic levels in each bay, we categorized the bays into stickleback- or perch-dominated bays. Earlier studies show an inverse relationship between stickleback and perch abundances (Bergström et al. 2015). We calculated a mesopredator index as the total number of sticklebacks in each bay divided by the total number of perch and stickleback together. Thus, if the bay had stickleback present but no perch, the index equalled 1; if the bay had perch present but no stickleback, the index equalled 0. The index revealed a reciprocal relationship between perch and stickleback: 27 of the 32 bays had a mesopredator index either higher than 0.80 or lower than 0.20, and no bay had an index between 0.40 and 0.60 (Fig. S1.4). Based on this, we assigned 16 bays with an index < 0.4 as perch-dominated and 16 bays with an index > 0.6 as stickleback-dominated.

We then examined whether increased relative abundance of stickleback versus perch may have changed the food web structure by comparing the relation between topographic openness (which promotes marine benthic production) and the quantity of organisms at different trophic levels in the two types of bays. According to the hypothesis of exploitation ecosystems (Oksanen et al. 1981, Oksanen and Oksanen 2000), the biomass of the first and third trophic level should increase with production in a system with three trophic levels, while the second and fourth trophic level should increase with production in systems with four trophic levels (see also Eriksson et al. 2012). This leads to the hypothesis that the cover of filamentous algae and stickleback abundance (1^st^ and 3^rd^ trophic level, respectively) should increase with increasing topographic openness in the stickleback-dominated bays, while only the crustacean macro-herbivores (2^nd^ trophic level) should increase with increasing topographic openness in the perch-dominated bays (Oksanen et al. 1981, Oksanen and Oksanen 2000, Eriksson et al. 2012). Accordingly, the CPUE of three-spined stickleback, the dry-mass of crustacean macro-herbivores, and the cover of filamentous algae were analyzed using General Linear Models (GLM) with topographic openness and bay type (fixed factor) as explanatory variables. Since we were specifically interested in the relationship between each of the trophic levels and topographic openness in bays dominated by perch or stickleback; different regression models were fitted to each bay type by nesting topographic openness (with bay type) (single call using the lm function in R 3.5.2; R Core Team, 2018). In contrast to a model with an interaction between the two bay types, which would test whether the slopes differ regardless of whether both are significant on their own or not; the nested model tests if there is a relationship between each trophic level and topographic openness for each bay type, independent of the relationship (or not) in the other bay type. All variables were normalized using log_10_ or square root transformations.

### Drivers of stickleback phenotype composition

We tested whether the relative abundance of three-spined stickleback phenotypes correlated with bay production and the abundance of predators (Fig. 1) by modelling the proportion of incompletely plated three-spined stickleback individuals in the bays, with a general linear model that included the explanatory variables topographic openness (Ea) and CPUE perch (# larger than 25 cm) (using the lm function in R 3.5.2; R Core Team, 2018). Since we only defined the stickleback as completely or incompletely plated, their relative proportions of the phenotypes are the inverse of each other. Therefore, we did not analyse the distribution of the completely plated phenotype separately. In the perch-dominated bays (n = 16) we found very few three-spined stickleback (mean=5.5, SD=5.3, max=22; CPUE). Thus, to get a reliable and comparable estimate of the fraction of plate phenotypes, we only included the stickleback-dominated bays (n = 16, Table S1), where we could collect ≥ 30 individuals. 492 individuals from the 16 stickleback-dominated bays were morphotyped and used for the analysis of plate phenotype distribution. Data was logit (proportion of fully plated individuals) and square root (Ea and # perch > 25 cm) transformed to fit assumptions of normality.

### Stickleback diet analyses

We used a metabarcoding approach to analyse diet differentiation between sympatric stickleback plate phenotypes. In many of the bays with few sticklebacks, we only found one of the two plate phenotypes. To avoid bias in stomach content analyses caused by the food availability in those bays with only one phenotype present, we randomly selected 177 individuals from 10 bays where both plate phenotypes were present (Table S1). In the laboratory, the stomachs were dissected and flushed with 80% EtOH to remove all stomach contents and stored at −20° C in 80% EtOH. DNA was then extracted from the stomach contents using the UltraClean Tissue and Cells DNA Isolation Kit (MO BIO Laboratories). The dual PCR amplification method was then used for Illumina MiSeq library preparation (Bourlat et al. 2016). The amplicon primers were based on Leray et al. (2013) yielding a 313 bp fragment targeting the cytochrome c oxidase subunit 1 mitochondrial gene (CO1), and a blocking primer for *G. aculeatus* was also used to prevent amplification of predator DNA (all primer sequences can be found in Jakubavičiūtė *et al*. (2017). Illumina MiSeq produced 30,103,790 paired-end reads of 300 bp in length. The processing steps were performed using Qiime (Quantitative Insights into Microbial Ecology) version 1.9.1 (Caporaso et al. 2010) and custom python scripts. Paired-end joining was done using the Qiime fastq-join tool. Primer sequences were removed using a custom python script, and remaining chimeric reads were excluded using UCHIME (Edgar et al. 2011). The Bayesian clustering algorithm CROP was used to cluster the sequences into operational taxonomic units (OTUs; Hao et al. 2011). Taxonomic assignment was performed with the Uclust software implemented in Qiime (Edgar 2010), using a 97 % similarity limit when comparing the CO1sequences with our own reference database of Chironomidae, Nemertea, Xenacoelomorpha and Oligochaeta, combined with barcodes of Swedish Echinodermata, Mollusca, Cnidaria and Arthropoda from the Swedish Barcode of Life database (SweBol). Detailed methods for DNA extraction, amplicon library preparation, and bioinformatic analyses of the current samples can be found in Jakubavičiūtė et al. (2017). The raw sequence data are available as fastq files from the NCBI sequence read archive under accession number SRP101702, BioProject number PRJNA378633.

For analysis, the OTU tables were converted to a presence/absence matrix. The prey species were divided into broad taxonomic groups relevant for the ecology of the system: crustacean mesograzers (amphipods, isopods, and mysids), molluscs (gastropods and bivalves), benthic worms (polychaetes and annelids), insects (chironomids and others), and zooplankton/meiofauna (cladocerans, copepods, and ostracods). For the complete list of prey taxa found in this study, see Table S1 in Jakubavičiūtė et al. (2017). A table of presence or absence of prey groups in the stomach of each fish was compiled. Differences in the occurrence of prey in the stomachs of completely compared to incompletely plated phenotypes, was analysed using the multivariate statistics package Vegan (version 2.5-6) in R. First, we performed an Analysis of Similarities (ANOSIM) to test if there was a difference in the composition of prey in the stomachs of completely and incompletely plated sticklebacks (model: species composition of prey ∼ plate phenotype). Then we plotted prey groups and the centroids of plate phenotypes using the information from an unconstrained NMDS ordination, to identify which prey groups contributed most to the differences between plate phenotypes. For each of the prey groups that was identified by the visual inspection (amphipods, mysids, bivalves and polychaetes; see results) we analysed the difference in stomach content between completely and incompletely plated fish, using a generalized mixed-effects model using the lme4 package in R (version 3.5.2; Bates et al. 2015). The model was based on the binomial error distribution with a logit link function (logistic regression), and included plate phenotype as a categorical explanatory variable, and bay as random factor.

### Comparing food web relations in the field with experimental data

To get an indication of the strength and causality of documented field patterns, we compared the field data with the results from a previous study that experimentally manipulated the abundance of larger predatory fish (Sieben et al. 2011b). This study was performed at the Askö Laboratory, Stockholm University, located within the area of the field study. Sieben et al. (2011b) excluded larger predatory fish in the field using cages (length: 1.2 m, width: 0.55 m, height: 1.0 m), covered with a net with a mesh size of 14 mm. This allowed sticklebacks, but not their predators, to enter the cages. Ten cages with a complete net (closed cages excluding all larger fish) and ten cages that had a 0.6 m^2^ opening in the net along one side of the cage (open cages that allowed larger fish to enter), were put in the water at ca 1 m depth from June 22^nd^ to September 17^th^ in 2008 (randomized block design). To create a large gradient in production, half of the cages in each treatment were continuously enriched with nitrogen and phosphorus using coated slow-release N-P-K fertiliser pellets (Plantacote Depot 6 M, Urania Agrochem, Hamburg, Germany). Stickleback abundances in the cages were monitored by snorkelling after one and two weeks. Crustacean macro-herbivores (amphipods and isopods) were sampled after 1 week and at the end of the experiment (after 2 months), using pre-weighed and cleaned bundles of bladderwrack that were put in each cage (41.7 ± 1.4 g DW, mean ± SE). The development of algae cover was estimated after four weeks by measuring the average thickness of filamentous algae (in cm) (mainly *Ectocarpus siliqulosus* and *Pylaiella littoralis*) that covered the bottom of each cage. Nitrogen concentrations (total-Nitrogen = NH_4_ + NO_-3_ + NO_-2_) of the water was sampled from each cage after one week and one month of experimental time. For further detail of experimental design and sampling procedures, see Sieben et al. (2011b).

We tested the general hypothesis that potential production and the abundance of predators determine food web structure, by comparing the relationship between nitrogen concentrations and the quantity of organisms at different trophic levels in the two types of cages (open cages with four trophic levels, and closed cages with only three trophic levels). To make the analyses comparable, we constructed identical General Linear Models (GLM) as for the analyses of the survey data on relations in different bay types (see section “*Analysing food web structure*” above). Accordingly, three-spined stickleback (average of all counts), crustacean macro-herbivores (average of ash-free dry-mass from July and September), and filamentous algae (thickness of cover in cm), were analyzed using General Linear Models (GLM) with nitrogen concentrations and cage type (fixed factor) as explanatory variables. Different regression models were fitted to each cage type by nesting nitrogen concentrations within cage type (single call using the lm function in R 3.5.2; R Core Team, 2018). All variables were normalized using log_10_ or square root transformations.

## Results

### Changes in predator-prey relations along the coast

Juvenile three-spined stickleback abundances increased strongly along the western coast of the central Baltic Sea between 1996 and 2017 (Fig. 2). The best fitting model only included year as an explanatory factor, disregarding the spatial sampling structure (exponential fit: F_1,20_=42.2, R^2^_adj_=0.66, p<0.001). In contrast, juveniles of the dominating coastal predator perch decreased linearly between 1996 and 2017 (linear fit 1996-2017: t=−2.3, p=0.034), and sampling depth was included as a very important factor in the best fitting model (abundance increased with sampling depth: t=4.1, p<0.001; Full model: F_2,17_=12.2, R^2^_adj_=0.54, p<0.001; Fig. 2). Thus, the relative abundance of juveniles of top- and mesopredators have changed dramatically over the past decades, likely reflecting a changed trophic structure along the coast.

**Figure 2:**
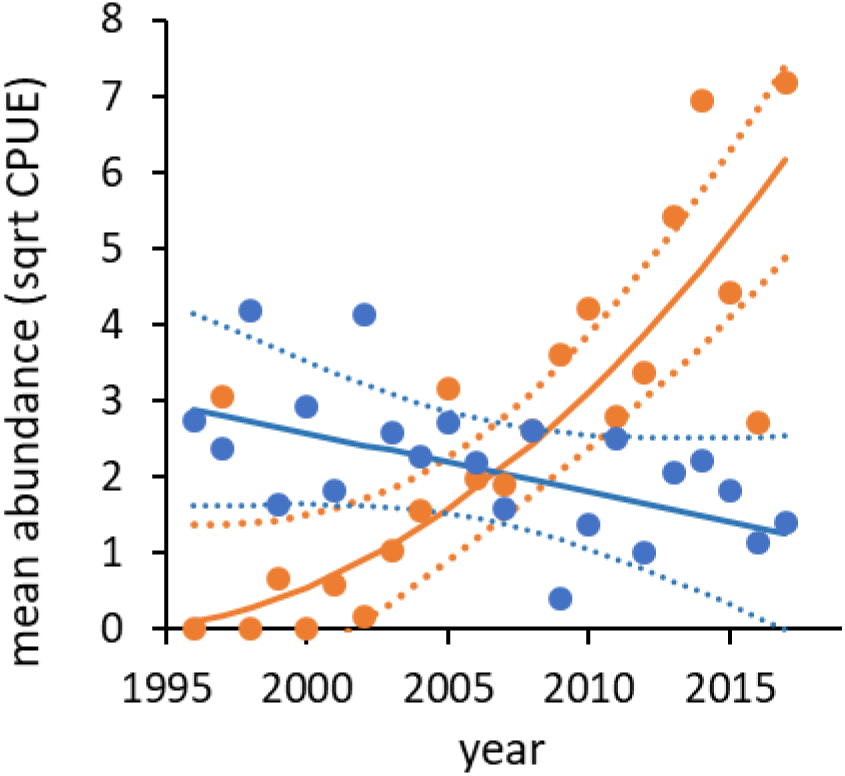
Temporal trends in three-spined stickleback (orange solid line) and perch (blue solid line) juvenile abundances (averages of square rooted numbers); between 1996 and 2017. For perch, the values are adjusted to account for differences in sampling depth. The solid lines show significant models describing the development over time, for perch it is partial residuals from a models accounting for sampling differences in depth. Dotted lines show 95% confidence interval around the predicted values.

### Differences in food-web dynamics along the coast

The food web field survey indicated the importance of production and predation in regulating the three-spined stickleback population; stickleback abundance increased with increasing topographic openness (Ea) and decreased with increasing abundance of large predatory fish (# perch > 25 cm; Model fit: R_adj_=0.43, F_1,29_=12.7, p<0.001). The variability in food web structure across the 32 bays supported the hypothesis that the dramatic shift from perch to three-spined stickleback has changed local food web dynamics, from a system characterized by four trophic levels to a system characterized by three (Fig. 3). Filamentous algae and stickleback (trophic levels 1 and 3) increased with topographic openness in bays dominated by stickleback (trend for algae: t=1.9, p=0.07; stickleback: t=3.8, p<0.001), but not in bays dominated by perch (algae: t=1.5, p=0.144: stickleback: t=0.0, p=0.999; full model algae: R_adj_=0.80, F_4,21_=26.6, p<0.001; full model stickleback: R_adj_=0.88, F_4,28_=59.5, p<0.001; Fig. 3). In contrast, mesograzer biomass (trophic level 2) increased with topographic openness in bays dominated by perch (t=2.9, p=0.007), but not in bays dominated by stickleback (t=0.0, p=0.986; full model: R_adj_=0.75, F_4,28_=25.2, p<0.001; Fig. 3).

**Figure 3:**
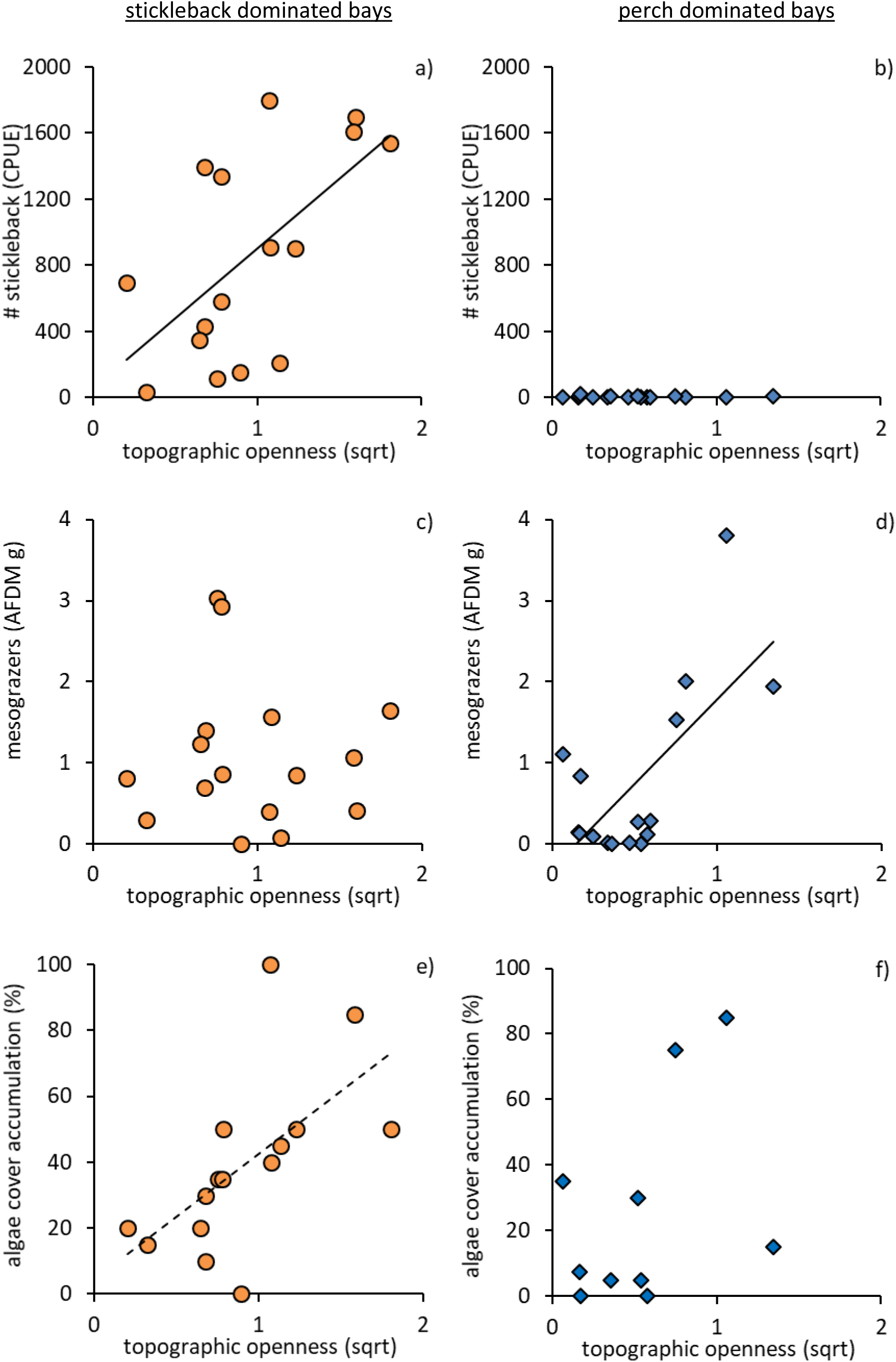
The relationships between different trophic levels and topographic openness (as a proxy for marine benthic production), in bays dominated by three-spined stickleback (a, c & e; orange circles) or perch (b, d & f; blue diamonds). a-b) the density of adult three-spined sticklebacks (3^rd^ trophic level), c-d) the biomass of crustacean macro-herbivores (2^nd^ trophic level), and e-f) the development of filamentous algae cover (1^st^ trophic level); Solid trend lines show significant relationships determined by linear regressions, dotted line show a trend towards significance (p=0.07). Note that the cover of filamentous algae are missing from eight bays (e-f) due to over-sedimentation of recruitment substrates.

Reanalysis of the Sieben et al. (2011b) experiment data supported the notion that changes in perch and stickleback abundances have changed food web dynamics along the coast (Fig. 4). In the closed cages with three trophic levels, filamentous algae and stickleback (trophic levels 1 and 3) increased with nitrogen concentration in the water (algae: t=3.7, p=0.002; stickleback: t=3.0, p=0.009), but not in open cages where perch could enter (algae: t=1.7, p=0.112: stickleback: t= −1.3, p=0.211; full model algae: R_adj_=0.42, F_3,16_=5.6, p=0.008; full model stickleback: R_adj_=0.32, F_3,16_=4.0, p=0.026; Fig. 4). In contrast, there was a trend for an increase in crustacean biomass (trophic level 2) with nitrogen concentration in the open (t=1.9, p=0.075), but not in the closed cages (t=1.2, p=0.244; full model: R_adj_=0.56, F_3,16_=9.1, p<0.001; Fig. 4). Amphipods, the most common crustacean mesograzer group, increased with nitrogen concentration in the open cages (t=2.2, p=0.040), but not in the closed ones (t=−0.0, p=0.966; full model: R_adj_=0.39, F_3,16_=5.0, p=0.012; Fig. 4c&d).

**Figure 4:**
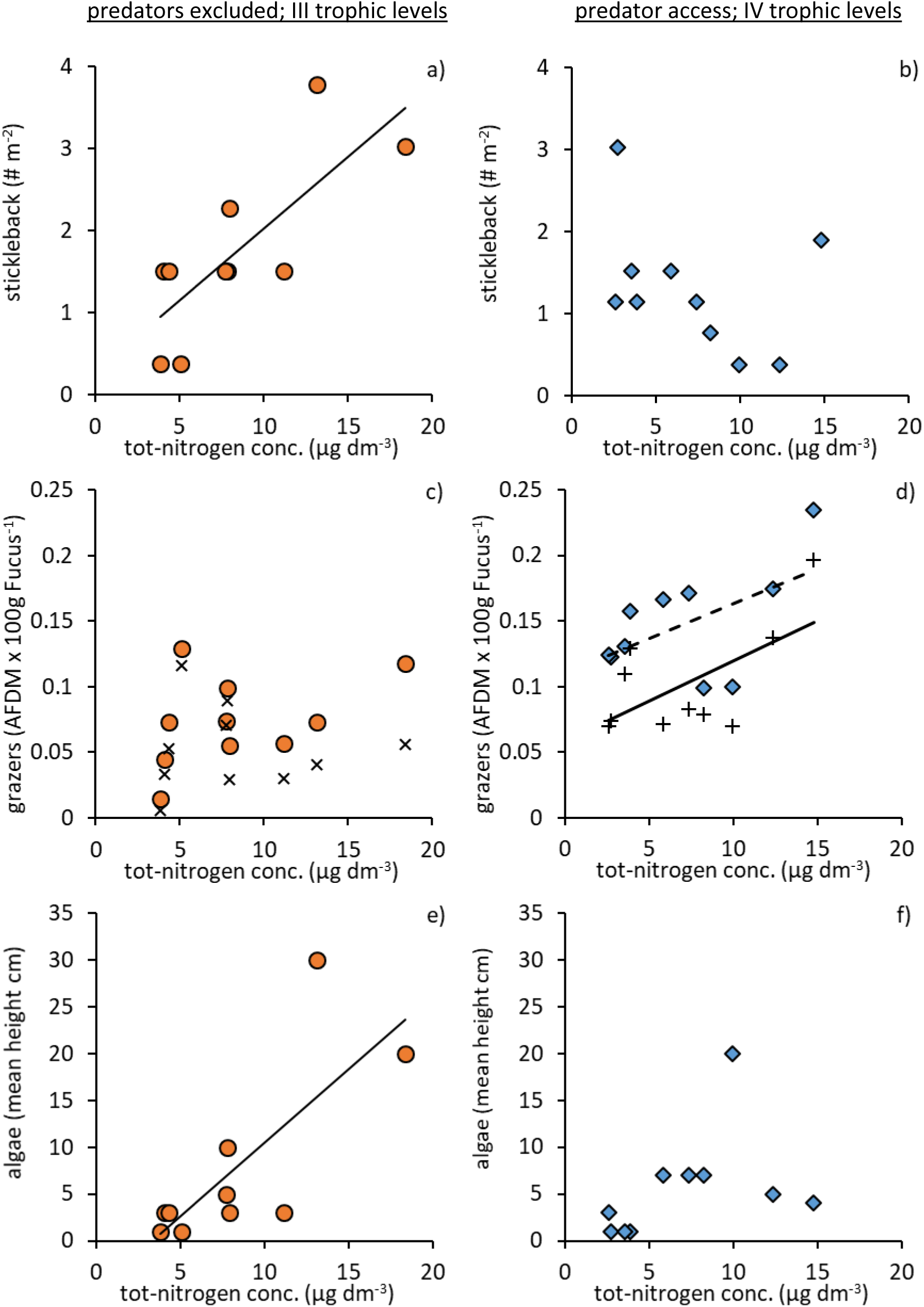
The relationships between different trophic levels and nitrogen concentrations in the water, in experimental cages with three (closed for larger predatory fish: a, c & e; orange circles) or four trophic levels (open for larger predatory fish dominated by perch: b, d & f; blue diamonds). a-b) the density of adult three-spined sticklebacks (3^rd^ trophic level), c-d) the biomass of crustacean macro-herbivores (2^nd^ trophic level), and e-f) the development of filamentous algae cover (1^st^ trophic level). In c and d, the x’s and +’s show the dominating macro-herbivore group amphipods. Solid trend lines show significant relationships determined by linear regressions, dotted line show a trend towards significance (p=0.075) for all crustacean macro-herbivores. Modified from Sieben et al. (2011b).

### The distribution of stickleback plate phenotypes along the coast

We observed two distinct stickleback phenotypes corresponding to completely (fully) and incompletely (partially/low) plated phenotypes and very few intermediates (Fig. 5). The proportion of incompletely plated individuals increased with topographic openness (t=2.7, n=16, p=0.019) and decreased with the abundance of perch larger than 25 cm (t=−2.5, n=16, p=0.026; Full model: R_adj_ = 0.39, F_2,13_=5.9, p=0.015, Fig. 6). This indicates a potential change in selection regime on plate phenotypes depending on productivity and predation. The incompletely plated phenotype was five times more common than the completely plated phenotype, and their relative abundance increased from 75% in the most enclosed bays to 90% in most open bays (Fig. 6a). At the same time, bays where perch was more abundant had a lower proportion of incompletely plated stickleback than expected from the topographic openness of the bay (Fig. 6b).

**Figure 5:**
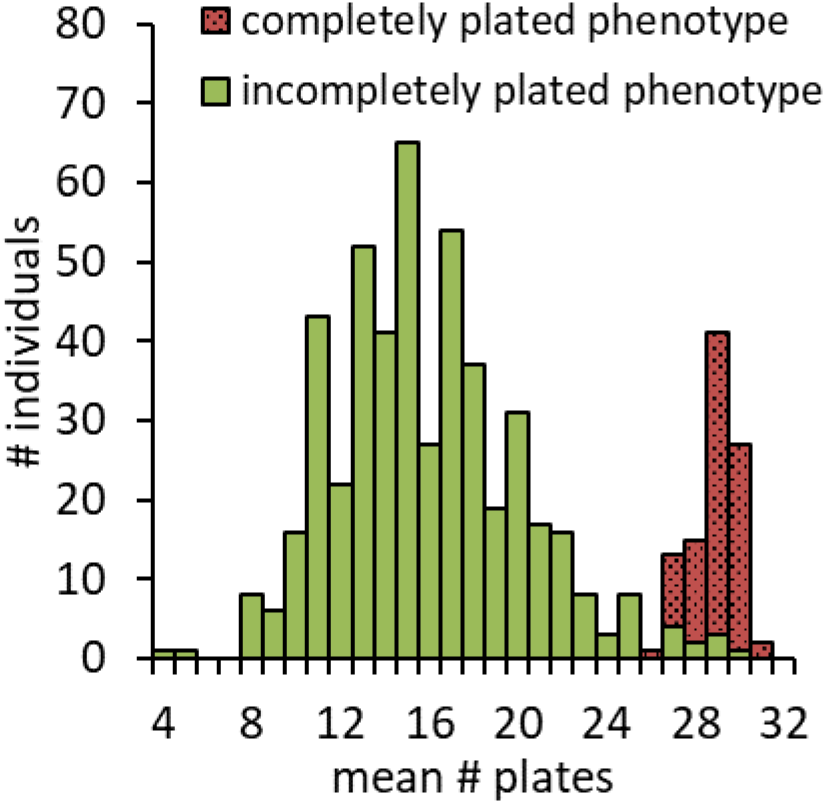
Frequency distribution of the average number of lateral plates, indicating the completely (fully) and incompletely (partial and low) plated three-spined stickleback phenotypes.

**Figure 6:**
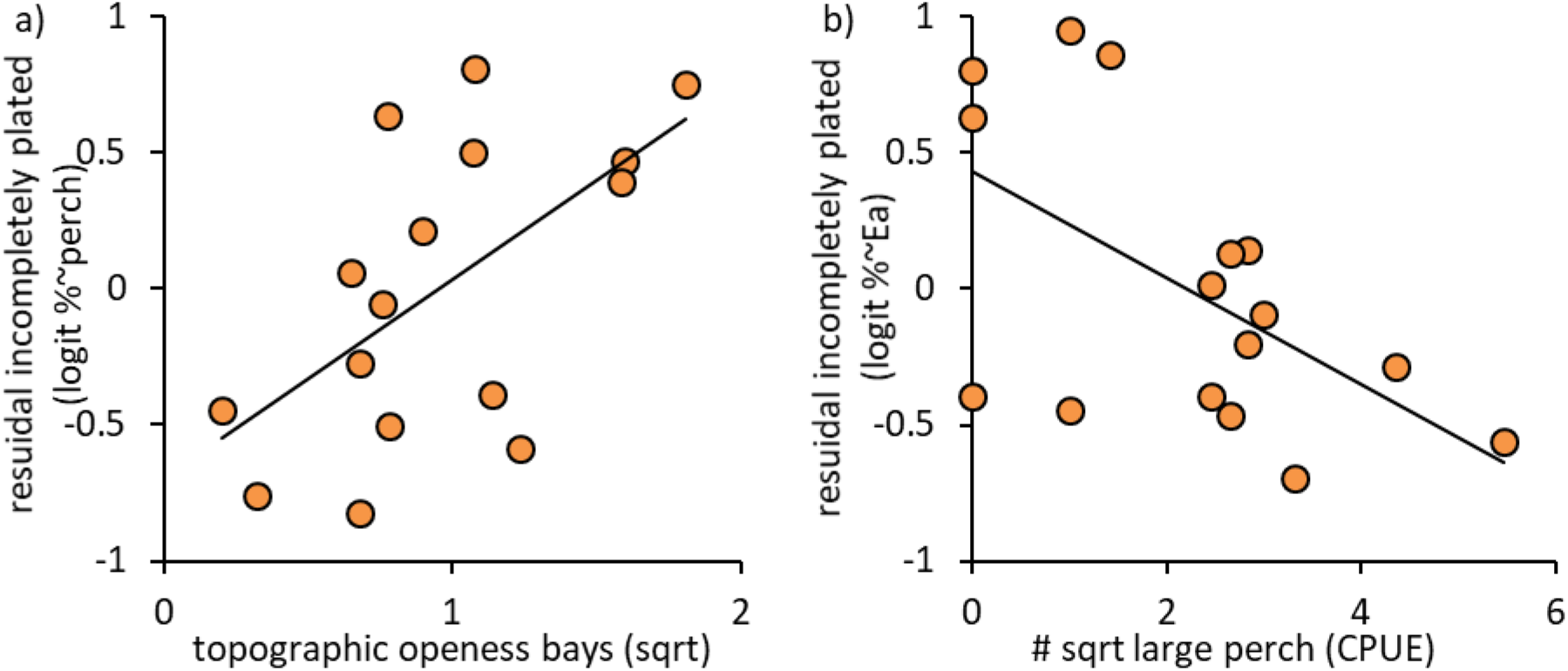
The residual variation of the proportion of incompletely plated adult sticklebacks (logit transformed data) explained by: a) topographic openness, after taking the abundance of large perch (>25 cm) into account; and b) the abundance of large perch after taking the topographic openness into account. Trend lines show significant relationships determined by linear regressions.

### Diet of the plate phenotypes

Metabarcoding of stickleback stomach content revealed significant differences in the diet of the two plate phenotypes (ANOSIM: R=0.075, p=0.037). The overall difference was small, because almost all sticklebacks had some insect (98.3 ±1.0%; chironomids; others) and/or zooplankton/meiofauna prey in their stomachs (99.4 ±0.5%; ostracods: 79.7±3.0%; cladocerans: 97.2±1.2%; copepods: 93.2±1.9%; mean±SE). However, the NDMS ordination highlighted four prey groups that contributed more strongly to variation along the ordination axes: amphipods, bivalves, mysids and polychaetes (Fig. S3). Analysing these groups separately, it was clear that the completely plated sticklebacks included amphipods more often in their diet compared to the incompletely plated ones (logistic regression: n=172, χ^2^=7.7, p=0.006; Benjamini Hochberg corrected p = 0.023). Amphipods are the dominant benthic herbivores in the area (Eriksson et al. 2009, Sieben et al. 2011b) and 56.1±7.8% of the fully plated individuals had amphipods in their stomachs, compared to 30.1±4.0% of the incompletely plated. Meanwhile, the proportion of occurrence of bivalves, mysids and polychaetes did not differ between stickleback phenotypes (logistic regression, n=172; bivalves: χ^2^=0.66, p=0.418; mysids χ^2^=0.19, p=0.662; polychaetes: χ^2^=0.17, p=0.683).

## Discussion

Management of restructured ecosystems poses a great challenge because emerging ecological interactions can change food web dynamics. Our results support the hypothesis that the increase of the three-spined stickleback in the Baltic Sea may have restructured coastal ecosystems and altered the existing trophic cascade, from a system characterized by four trophic levels to a system characterized by three trophic levels (supporting the hypothesis of a changed food web structure outlined in Fig. 1). In bays dominated by the three-spined stickleback, organism biomass at the first and third trophic levels (filamentous algae and stickleback) increased with topographic openness (which increases benthic production), while in bays dominated by the larger predatory fish perch, only biomass at the second trophic level (herbivores) increased with topographic openness. These patterns were supported by reanalysis of Sieben et al.’s (2011b) experimental data, confirming that declining predator abundances interacts with primary production to cause a trophic cascade along the coast. In spite of high variation leading to weak statistical models, the experimental results were remarkably similar to what was observed in the field. Our analyses of coastal time-series data show that the increase in three-spined stickleback abundance was followed by a decline in the abundance of juveniles of its main predator perch between 1995 and 2005. Suggested causes for the decline of perch include loss of recruitment habitats (Sundblad and Bergström 2014), but recent studies indicate that sticklebacks can also control the recruitment of perch by feeding on its juvenile stages (Byström et al. 2015, Eklöf in review).

In addition to the predicted ecological effects (Oksanen et al. 1981, Nisbet et al. 1997, Oksanen and Oksanen 2000, Sieben et al. 2011b), our results indicate the possibility for density-dependent selection and eco-evolutionary feedbacks in the areas now dominated by the three-spined stickleback. In stickleback-dominated bays, the ratio of incompletely *vs* completely plated armour phenotypes increased with topographic openness, and decreased with the abundance of large predatory perch. Moreover, the two phenotypes had small but significant differences in their diets. Specifically, more completely plated than incompletely plated stickleback ate amphipods; the key benthic mesograzers in the areas (Donadi et al. 2017). This suggests that a recent shift in stickleback selection regime across the Baltic Sea – from high predation pressure to high intra-specific competition – may alter the effects of sticklebacks on food web dynamics (supporting the hypothesis of an eco-evolutionary feedback outlined in Fig. 1b).

Low abundance of large predatory fish was associated with high abundance of the incompletely plated phenotype. This is interesting since the three-spined stickleback has, since the last glaciation, repeatedly colonized and adapted to predator-free freshwater systems, by evolving a low plated phenotype characterised by a reduced number of bony lateral plates (Bell 2001, Leinonen et al. 2011a). This striking example of repeated evolution is founded in a consistent reuse of globally shared standing genetic variation (Schluter and Conte 2009, Jones et al. 2012). The lateral plate number is a trait under selection (Barrett et al. 2008) and ca 75 % of the variation is determined by a major locus, ectodysplasin (EDA; Colosimo et al. 2004, Le Rouzic et al. 2011). Plates act as armour against gape limited toothed predators (Reimchen 1992, Bergström 2002) and there is both observational and experimental evidence to support the notion that the low plate phenotype is favoured in freshwater populations due to reduced predation by fish (Bell et al. 2004, Cresko et al. 2004, Kitano et al. 2008, Leinonen et al. 2012). Therefore, it is possible that the observed differences in distribution of the different plate phenotypes along the coast in fact reflects a temporal change in predation pressure due to declining predatory fish abundances.

Although the different stickleback phenotypes co-occur in the Baltic Sea (Jakubavičiūtė et al. 2018), our results indicate that they differ in their diets at least in coastal areas. The diet differences were caused by one prey group only: completely plated sticklebacks more often consumed amphipods. Amphipods are dominant benthic herbivores in the area (Eriksson et al. 2009, Sieben et al. 2011b), and they form a key grazer group that mediate cascading ecosystem effects in coastal areas across the world (Duffy and Hay 2000, Råberg and Kautsky 2007, Moksnes et al. 2008, Duffy et al. 2015, Donadi et al. 2017). Experimental testing or more controlled sampling of different phenotypes, while controlling for prey availability, is needed to evaluate how important this diet difference is for food web dynamics. However, it is known that changes in the distribution of stickleback plate phenotypes can have strong effects on ecosystem function in a number of ways (Harmon et al. 2009). Effects of rapid diversification of partially plated stickleback include changes in prey community structure, biomass and abundance (Matthews et al. 2016, Rudman and Schluter 2016, Schmid et al. 2019); elemental stoichiometry (El-Sabaawi et al. 2016), phosphorus excretion (Paccard et al. 2018) and phosphorus concentrations in the water (Matthews et al. 2016). In addition, adaptive shifts in mean trait values may lead to increased stickleback abundances, which in turn should change their ecological effects (indirect eco-evolutionary effects: Hendry 2017). To understand if similar effects are occurring in the Baltic Sea, potentially causing eco-evolutionary feedbacks that change the effects of the mesopredator release, experimental studies would be needed.

The relative contributions of different factors to the increase in three-spine stickleback abundance in the Baltic Sea are currently not known. Our results show that juveniles of the main coastal predator, perch, have decreased over the past decades, and that the selection for anti-predator traits (plating) may have become relaxed as the incomplete phenotype dominates the studied stickleback population. Yet, the causality needs to be established in future studies, as there are other factors that may contribute to the increased stickleback abundance. In particular, increasing temperatures and habitat change (Eriksson et al. 2011, Lefebure et al. 2014, Nilsson et al. 2014, Sundblad and Bergstrom 2014, Hansen et al. 2019). In fact, similarly to the Baltic Sea, the White Sea has shown an increase in three-spined stickleback abundance since the late 1990’s (Lajus et al. 2020). The increase in the White Sea has been attributed mainly to increased water temperatures and subsequent changes in habitat availability (Rybkina et al. 2017, Lajus et al. 2020).

In conclusion, the results indicate that a mesopredator release in the Baltic Sea has restructured of local food webs, and that eco-evolutionary effects may be involved. In the past decades, the mesopredatory three-spined stickleback has increased greatly in the western coasts of the central Baltic Sea, coinciding with local declines in juveniles of the important large predatory fish, perch (Eriksson et al. 2011, Olsson et al. 2019). Today the three-spined stickleback dominates a large proportion of bays along the Swedish Baltic Sea coast, especially those with a high topographic openness and high productivity. In stickleback-dominated bays, filamentous algae and stickleback increase in abundance when benthic production increases, but grazers do not exhibit a similar response – a clear indication of a mesopredator-dominated ecosystem (Oksanen and Oksanen 2000). At the same time, an increased proportion of incompletely plated individuals together with higher stickleback numbers, indicate a reduced selection for anti-predator traits and increased density dependent selection in bays with low abundances of perch and high benthic production. We also found small but significant differences in diets between the phenotypes, where the incompletely plated individuals eat less of the key benthic herbivore amphipods. These results, together with evidence for that changes in the trait distribution in three-spined sticklebacks can have variety of community and ecosystem effects (e.g. Matthews et al. 2016, Rudman and Schluter 2016, El-Sabaawi et al. 2016, Schmid et al. 2019, Paccard et al. 2018), indicate the potential for eco-evolutionary effects in the broad sense (de Meester et al. 2019); where ecological effects generate changes in selection regimes, which in turn feed back on a variety of ecological processes. However, to provide solid empirical predictions of eco-evolutionary consequences of the shift in selection pressures, we need experimental testing of the ecological effects of different phenotypes, behavioural studies to assess morphology-dependent mating patterns, and genetic studies to identify signatures of selection. Our current results emphasize the need to better understand how ecological feedbacks can change selective regimes, potentially driving rapid adaptation, and the need to take these effects into account in ecosystem management strategies (de Meester et al. 2019, Hendry 2019).

## Acknowledgements

We thank the property and fishing right owners around each of the 32 study bays for facilitating the fieldwork and a number of students, volunteers, and fish biologists for assistance in the field and laboratory: E. Anderberg, F. Ek, G. Johansson, P. Jacobson, C. Jönander, E. Mörk, L. Näslund, O. Pettersson, G. Lilliesköld Sjöö, S. Skoglund, M. van Regteren, M. van der Snoek, V. Thunell and L. Vik. The authors declare no conflict of interest.

