## Supplement 1 for "The rise of the three-spined stickleback – eco-evolutionary consequences of a mesopredator release"

### Supplement 1: Extended methods and results for sampling the food-web structure in bays along the west coast of the Baltic Sea

The distribution of three-spined stickleback ecotypes and the structure of the coastal food-web was analyzed by sampling the shallow-water fish community, invertebrates, vegetation and macroalgae in 32 bays along a 360 km stretch of the central Baltic Sea coast, in spring 2014 (Fig. S1.1, Table S1). The sites were distributed throughout the area to achieve variation in both abiotic and biotic factors, such as salinity, exposure, fish community structure and densities of different species. The investigated bays were chosen using the following criteria:

- Distance between bays was set to a minimum of 10 km unless they were separated by naturally occurring barriers (e.g. large bodies of open water or land areas) to assure sampling fish from different populations.
- Gradient in wave exposure (moderately exposed to extremely sheltered)
- Gradient in topographic openness (open to semi-enclosed bays)
- Gradient in salinity (4-7 practical salinity units (PSU))
- Sufficient areas with a depth between 0.5-3m
- Suitable nursery areas for three-spine stickleback and pike and perch (Sundblad et al. 2011, Nyström Sandman A. et al. 2013)

Fish were caught using Nordic multimesh gillnets. There is a standardized survey method in the European Union for this gear type (EN 14757:2005), which gives an estimation of the relative fish abundance and fish species distribution of the entire fish community in an area. The method is also used in the Swedish national fish monitoring program (Havs och vattenmyndigheten 2014). The Nordic gillnets were 30 meters in length, 1.5 meters in height and consists of 12 panels, each 2.5 meter in length with different mesh sizes in the following order: 43, 19.5, 6.25, 10, 55, 8, 12.5, 24, 15.5, 5, 35 and 29 mm knot-to-knot. Four nets were set in each bay at a depth between 1.5-3 meters >30 meters from each other, and >10 meters from both land and bay opening. The nets were placed between 16:00-19:00 and lifted between 07:00-09:00 the next morning. Depth of the net placement was measured at the two ends of each net when setting the net, while salinity, turbidity and time were recorded both when setting and lifting the nets. Turbidity was measured with a fluorometer (Aquafluor), salinity and temperature with a Multi 340i voltmeter (WTW, Germany).

All fish caught were identified to species, measured (total length, closest cm) and counted. The number of individuals of each fish species caught in all four nets were pooled and expressed as catch per unit effort (CPUE). CPUE was then used as an estimate of the fish densities within each bay. In each bay, we randomly collected 30 three-spined sticklebacks from the total amount of stickleback caught in the nets and stored them in plastic vials with ethanol (analytical grade 95 %). If less than 30 was caught in a bay, all individuals were collected.

Vegetation cover, the biomass of epifauna, zooplankton abundance, temperature and salinity, were measured at three to six randomly distributed stations; the number of stations depended on the size of the bay (Donadi et al. 2017). A station was defined as a 5 m-diameter area around a pre-selected way-point (marked with a buoy) at a depth of 0.5–3 m. Each station was placed at least 30 m apart. The percentage cover of macrophytes on the bottom (vegetation cover) was visually estimated by a snorkeler in three randomly placed 0.5 x 0.5 m frames, within the large area Fig. S1.2a). Epifauna was collected by a snorkeler in a randomly placed 0.2 x 0.2 m frame connected to a 1 mm-mesh bag (Fig. S1.2b). The randomization of frames was accomplished by the snorkeler, by closing her/his eyes, diving down in the water column and placing the frame blindly on the bottom (Fig. S1.2c). After identification in the laboratory, the biomass of the epifauna was estimated as gram ash-free dry mass using taxonspecific length : AFDM correlations. Zooplankton was collected using Apstein nets: a conical net with a mesh size of 80 µm connected to a circular frame with an opening of 25 cm. 3-5 vertical hauls were taken at each station: in deeper water 3 vertical hauls were taken and in shallower water 5, to make water volume more comparable. The collected samples were immediately fixed in formaline. Zooplankton was later identified to taxon level in the in the laboratory. Temperature and salinity of the water was measured using a multi-measurer (Multi 340i voltmeter, WTW, Germany).

The development of macroalgal cover was estimated using algal settling plates. A settlement plate consisted of a 5 x 5 cm unglazed ceramic tile. In each bay we placed two clay bricks with two tiles each (total area 1 $dm^2$) glued to the surface, at ca 1.5 m depth. To avoid shading and over-sedimentation, we placed the bricks in areas devoid of vegetation and on as hard substrate as possible (Fig. S1.2d). The bricks were placed in the water by snorkeling during the fish sampling, and extracted from each bay three months later in August 2014. The tiles where brought to the lab and total accumulation of filamentous macroalgal cover over the summer was determined by overlaying a 5 x 5 cm plexiglas sheet with 25 random dots and counting how many dots that covered an algae. Determining algal cover accumulation over a longer time period on settlement plates, is a good way to compare net production of algae between different experimental units in the Baltic Sea (e.g. Eriksson et al. 2006, Eriksson et al. 2009, Eriksson et al. 2012). However, it is a relative measure that can only be compared within a study. Filamentous macroalgae respond to nutrient enrichment by fast accumulation of biomass and are grazed on by small invertebrate mesograzers (Worm et al. 2002, Råberg and Kautsky 2007). Mesograzers are in turn eaten by fish and play a key role in the trophic cascades earlier demonstrated in the system (Eriksson et al. 2009, Donadi et al. 2017). The relative development of filamentous algae is a therefore good indicator of top-down grazer control or bottom-up resource control of primary biomass in the system.

We characterized the physical environment of the bays by estimating topographic openness, wave exposure and distance to the open sea. Bay topographic openness (Ea) is defined as Ea = 100 x At/a: where At is the cross sectional area of the bay opening, calculated by measuring depth and width of the smallest opening of the bay towards the sea (depth was measured in the field, width using satellite images and GIS methods); and a is the bay surface area estimated from satellite images using GIS methods. Wave exposure was estimated for each bay using a standard wave model based on fetch, topography and wind conditions (Buschbaum et al. 2016). Distance from the open sea was calculated by estimating the shortest water distance (m) from the opening of each bay to the baseline (the starting point for defining territorial sea). The baseline is calculated by drawing the shortest distance in a straight line between the outermost islands in a nautical chart.

Other explanatory variables related to nutrient availability and plankton production (N and P concentrations, turbidity and fluorescence of the water) was also recorded at each station, (see Donadi et al. 2017 for details on sampling).

Table S1. Location of 32 bays sampled across the extensive archipelago in the central part of the western Baltic Sea, Sweden, and the selection of bays for the different analyses. Plate form analyses only included stickleback dominated bays where 30 or more 3-spine sticklebacks were found and morphotyped. Diet analyses only included bays where we had metabarcoding data from both plate genotypes (fully or partially plated genotypes).

| Bay abbre-viation | Bay name (total 32 bays) | Lat-itude | Long-itude | Stickleback or Perch dominated bay | Plate-form analyses (16 bays) | Diet analyses (10 bays) |
|---|---|---|---|---|---|---|
| HATT | Hatten | 60.382 | 18.249 | perch | NO | NO |
| SAHO | Sandå Holme | 60.360 | 18.560 | perch | NO | NO |
| MAST | Lilla Måstensfjärden | 60.36 | 18.56 | perch | NO | NO |
| JARS | Järsön | 60.309 | 18.473 | stickleback | YES | NO |
| SFJA | Söderfjärden | 60.27 | 18.628 | perch | NO | NO |
| ENHO | Enholmarna - sundet Saltö | 60.270 | 18.628 | perch | NO | NO |
| BRAN | Brännskär | 59.693 | 19.148 | stickleback | YES | NO |
| SJAL | Långviken, Själbottna | 59.565 | 18.804 | stickleback | YES | YES |
| BYVI | Byviken | 59.392 | 18.657 | perch | NO | NO |
| ALGK | Älgkilen | 59.388 | 18.937 | perch | NO | NO |
| BOLV | Bolviken | 59.383 | 18.53 | perch | NO | YES |
| LOVO | Lövösundet | 59.272 | 18.675 | perch | NO | NO |
| KRYS | Kryssviken | 59.272 | 18.675 | stickleback | YES | YES |
| TORV | Torviken | 59.113 | 18.479 | perch | NO | YES |
| HASL | Hasslinge-Skäret | 59.105 | 18.243 | stickleback | YES | NO |
| HANS | Hansviken | 59.028 | 18.023 | perch | NO | YES |
| VARN | Värnöfladen | 59.015 | 18.384 | stickleback | YES | YES |
| SORV | Sörviken, Fifång | 58.841 | 17.716 | perch | NO | NO |
| BOCK | Bockholms-sundet | 58.829 | 17.616 | stickleback | YES | NO |
| LACK | Hamnhamn, Lacka | 58.747 | 17.572 | stickleback | YES | NO |

| | | | | | | |
|---|---|---|---|---|---|---|
| OLER | Östra Lermaren | 58.747 | 17.572 | perch | NO | NO |
| OSVI | Östra viken | 58.729 | 17.19 | stickleback | YES | YES |
| KUGG | Kuggviken | 58.728 | 17.444 | perch | NO | NO |
| BETN | Beten | 58.645 | 17.155 | stickleback | YES | YES |
| LBRE | Lilla Brevik | 58.644 | 17.021 | stickleback | YES | YES |
| OSTA | Östantillsviken | 58.409 | 16.892 | perch | NO | NO |
| BROT | Brottskären | 58.347 | 16.94 | stickleback | YES | NO |
| SORS | Sörsundsviken | 58.252 | 16.88 | stickleback | YES | NO |
| SKRA | Skräckskärs-klubben | 58.167 | 16.978 | stickleback | YES | NO |
| BANK | Bankeböte | 58.107 | 16.733 | stickleback | YES | YES |
| HEMV | Hemviken Ängsö | 58.004 | 16.719 | perch | NO | NO |
| HUML | Norra Hummeldalen | 57.949 | 16.79 | stickleback | YES | YES |

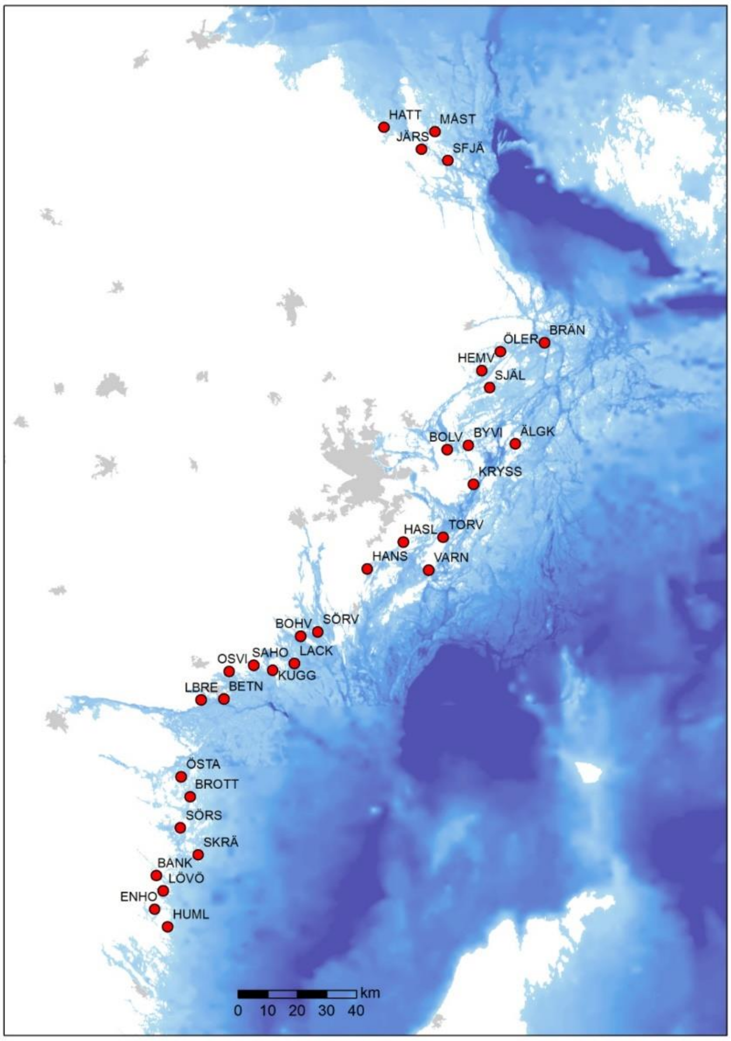


*Figure S1.1. The 32 bays sampled in 2014 along the extensive archipelago in the central part of the western Baltic Sea, Sweden.*

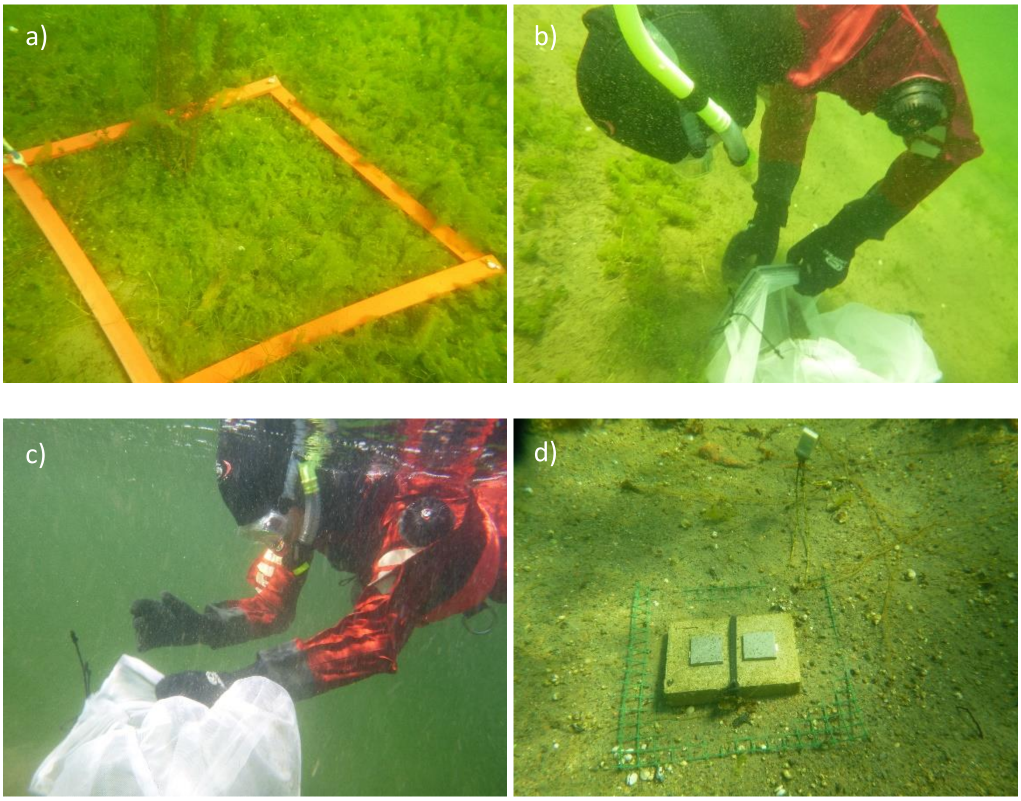


*Figure S1.2. Sampling a) vegetation cover and b) biomass of vegetation and epifauna, using quadrats randomly placed on the bottom at 0.5–3 m depth, within a 5 m-diameter area around each pre-selected way-point. c) snorkeler closing her eyes and diving to randomly place quantitative quadrate. d) settling plates glued a brick, placed on the bottom on a net to minimize overgrowing vegetation. Temperature logger is attached to the brick.*

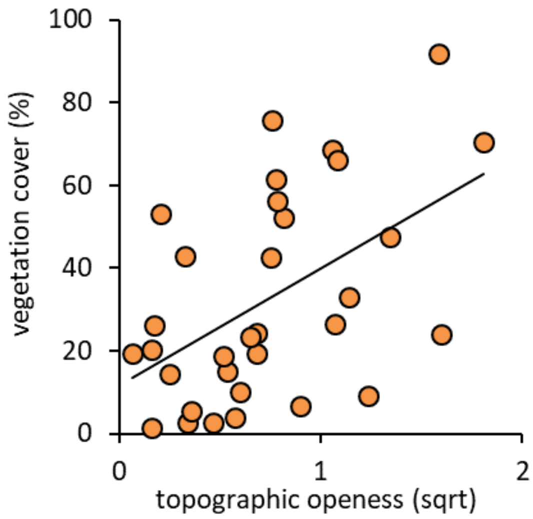


*Figure S1.3. The relationship between vegetation cover topographic openness in 32 bays; showing that topographic openness is a proxy for general community production. The trend lines show a significant relationships determined by a linear regression; r=0.50, t=3.2, p=003).*

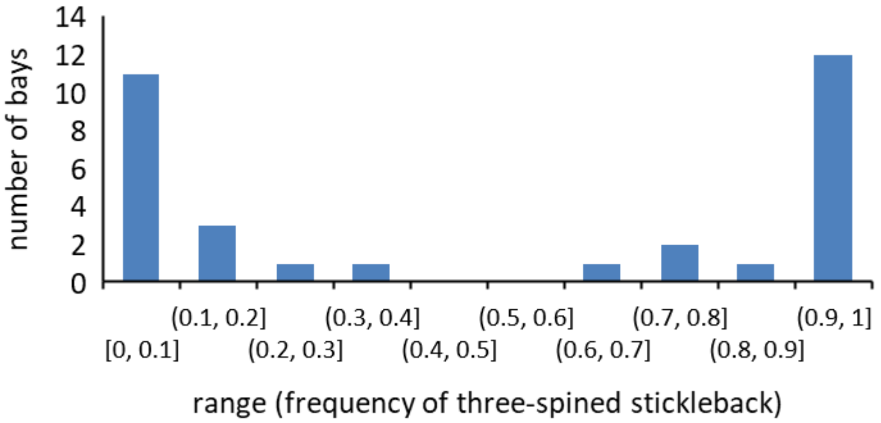


*Figure S1.4. Histogram showing the number of bays with different frequency of three-spined stickleback compared to perch. If the bay had stickleback present but no perch, the frequency had the value 1; and if the bay had perch present but no stickleback, the frequency had the value of 0.*
