## Supplement 2 for "The rise of the three-spined stickleback – eco-evolutionary consequences of a mesopredator release"

### Supplement 2: Analysing the distribution of three-spine stickleback plate genotypes

#### Methods

*Morphological determination of 3-spined stickleback*

In total, we determined the plate genotype (partial or full plating) for 560 three-spined stickleback individuals by measuring morphological characteristics. Eleven length parameters were documented from all individuals to the nearest millimeter using a Vernier caliper, according to Jones et al. (Jones et al. 2012): (1) total length; (2) standard length; (3) length of first spine; (4) length of second spine; (5) length of third spine; (6) length of caudal peduncle; (7) depth of caudal peduncle; (8) length of pelvic girdle; (9) head length; (10) width of pectoral fin base; (11) length of upper lip. Then the number of lateral plates were counted on both left and right sides to produce average number of later plates for each individual. The method for counting plates was that used by Aneer (1974). Plates were counted starting immediately after the operculum to the end of the caudal peduncle. Three-spined stickleback have three types of plates: plates near the operculum are typically circular, dermal formations which are not attached to any other body structure; plates along the sides of the body are typically oval or rectangular and are also not attached to other body structures; and 3) caudal plates vary in shape from circular to square to diamond-shaped and can be unattached to other body structures or can be the base of caudal scutes. We determined the plate phenotype by looking at gaps in the plate armour: partially plated fish had a clear gap between the main body plates and the caudal plates (the width of at least 2 body plates); fully plated fish had no clear gap in the plating. The entire length of the body of the fish is examined for presence of plates even if a gap is present to ensure plates are not missed. In total we morphotypes 560 individuals. The distribution of number of lateral plates overlapped slightly between the fully and partially plate phenotypes, but showed a clearly bimodal distribution with peaks at 15 and 29 plates for partially and fully plated individuals, respectively. We found no low plated individuals.

*Analyzing the distribution of plate genotypes*

The fraction of partially plated 3-spined stickleback individuals in the bays were analysed using a General Linear Mixed Model (GLMM) including explanatory variables known to determine the abundance of stickleback in the area (same as for CPUE in supplement 2: see Donadi et al. 2017); meaning that the model included explanatory variables related to topographical openness (physical parameter), zooplankton and epifauna abundance (food availability), and # perch larger than 25 cm (predation). The fraction was calculated as a log-response ratio; calculating the natural logarithm of the ratio in each bay (fraction=LN(# partially plated/# fully plated)). In half of the bays we found very few three-spined sticklebacks, meaning that the fraction was calculated on two to three individuals only. Thus, to get a reliable and comparable estimate of the fraction, we only included into the analyses those 16 stickleback dominated bays where we were able to collect ≥ 30 individuals (Table S1).
